# Dosa: A method to covalently barcode proteins for high throughput biochemistry

**DOI:** 10.1101/2025.09.21.677650

**Authors:** Yashwanth Ashok, Kerry L. Bubb, Curran Oi, Sayeh Gorjifard, Josh T. Cuperus, Christine Queitsch, Stanley Fields

## Abstract

Deep mutational scanning couples a protein’s activity to DNA sequencing for high throughput assessment of the effects of all single amino acid substitutions, but it largely uses indirect assays, like growth, as proxy for protein activity. Here, we covalently link variant proteins *in vivo* to an RNA barcode by fusing them to *E. coli* tRNA (m_5_U_54_) methyltransferase TrmA (E358Q), which forms a covalent bond with a tRNA stem-loop. Following cell lysis, variant proteins are separated *in vitro* according to their biochemical properties and identified by their barcodes. We use this method, Dosa, to analyze a large pool of FLAG epitope variants for binding to an anti-FLAG antibody, to profile the cleavage preferences of variants of enteropeptidase and human rhinovirus 3C protease, and to measure the solubility of several hundred Aβ(1-42) variants. This method should be amenable to numerous biochemical assays with proteins produced in *E. coli* or mammalian cells.

## Introduction

Mutations can affect fundamental properties of proteins, such as their solubility, abundance, stability or activity. The assessment of these properties is critical for understanding of a protein’s function and determining whether a mutation in a human protein leads to a pathogenic phenotype. However, traditional biochemical approaches that rely on the purification of individual variant proteins and a comparison of their properties to the wild type are costly and time-consuming, and cannot even remotely be applied to the expanding and vast number of known human protein variants. Thus, deep mutational scanning approaches^1^ are increasingly being applied. These analyze a library encoding all single amino acid substitutions in a protein, a selection for the protein’s activity, and DNA sequencing to determine the enrichment of each protein variant in the selection. Such approaches typically use cellular assays such as growth selections and flow sorting^2–4^ and largely indirect readouts, with growth or fluorescence often serving as a proxy for biochemical activity. For example, the aggregation of thousands of amyloid-β variants was assessed using a growth assay in yeast^5^. A direct measurement of protein properties such as aggregation, stability and solubility does not fit readily into the deep mutational scanning framework.

Efforts to more directly link genotype to phenotype rely on protein display methods. *In vitro* protein display methods like mRNA display, in which a puromycin moiety covalently links an mRNA to the protein it encodes, preserve the linkage even if the proteins are subject to harsh conditions, such as a denaturant or heat^6,7^. However, these methods require proteins to fold in the absence of a cellular environment, thereby failing to leverage protein quality control systems such as chaperones and post-translational modifications. *In vivo* display methods like cell surface display^8,9^ employ bacteria, yeast, mammalian or other cells to express proteins on their surface. But cells are fragile and can lose the genotype–phenotype linkage under harsh conditions.

Protein display methods can combine *in vivo* expression with *in vitro* assays. They couple a nucleic acid-binding domain such as a transcription factor^10,11^ or phage MS2 coat protein, as in *in vivo* tethered function assays^12^ and *in vivo* mRNA display,^13^ which can assay subcellular localization and protein interactions. In an approach called CasPlay, fusion proteins with a catalytically dead version of Cas9 (dCas9) were linked *in vivo* to their corresponding guide RNAs and used to evaluate *in vitro* peptide-antibody binding^14^. Label-seq, an improvement on *in vivo* mRNA display, linked fusion proteins *in vivo* to circular RNA barcodes and measured the abundances of the variants after cell lysis^15^. However, these linkages are noncovalent and thus are not usable in biochemical assays that employ elevated temperature or denaturing conditions.

We sought to develop an *in vivo* protein display method that combines the advantages of having cells produce properly folded proteins and covalently linking them to RNA barcodes. Our method, Dosa, relies on *E. coli* TrmA, a 42 kDa protein that methylates uracil-54 in the T-arm of tRNA, a nearly universal modification on tRNA. The TrmA (E358Q) mutant makes a covalent bond to tRNA at uracil-54, with a 19-nucleotide stem-loop region of the tRNA T-arm sufficient for the bond to occur^16^. We show that TrmA (E358Q) barcoded fusion proteins can be subjected to assays for protein binding, protease cleavage and solubility and their activities scored via counts of DNA sequencing reads.

## Results

### Development of an *in vivo* barcoding strategy

We leveraged the ability of the TrmA (E358Q) mutant to link covalently to tRNA in vivo to develop the Dosa method. In Dosa, the protein of interest is expressed from a plasmid as a fusion protein with *E. coli* TrmA (E358Q) (Fig. 1A). From the same plasmid backbone, we also express an RNA composed of the 19-nucleotide stem-loop TrmA-binding site of the tRNA T-arm, a unique 10-base barcode sequence and flanking sequences to act as PCR priming sites. Upon co-expression of protein and RNA, we anticipated that the barcode RNA would covalently link in *E. coli* cells via the RNA stem-loop to the TrmA (E358Q) fusion. Each cell acts as a barrier to prevent barcodes expressed in one cell from linking to a non-cognate TrmA fusion protein expressed in a different cell. We expressed wild-type (WT) and mutant TrmA proteins in *E. coli* and observed that TrmA (E358Q) covalently linked to endogenous tRNA whereas WT TrmA and the catalytically inactive TrmA (E358Q, C324A)^17^ did not (Fig. S1).

**Fig. 1.**
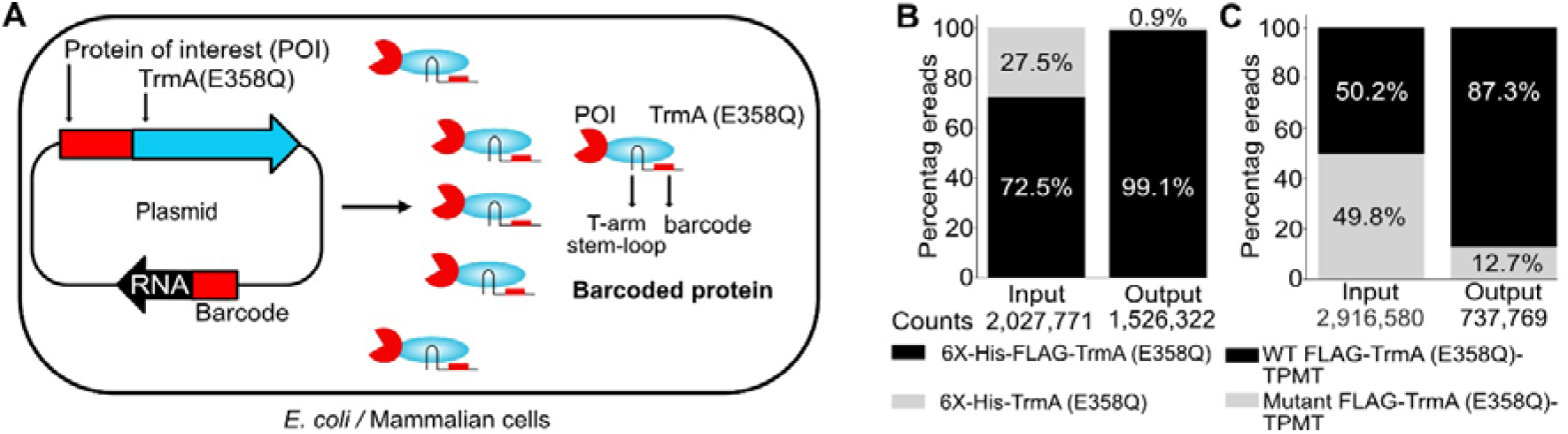
Development of the Dosa approach to barcode proteins *in vivo*. **A**. Overall scheme of barcoding proteins. A single plasmid encodes TrmA (E358Q) as a fusion protein with a protein of interest (POI). A separate transcript encodes the tRNA T-arm stem-loop (which binds to TrmA) and a barcode sequence flanked by PCR priming sites. Expression of the TrmA (E358Q) construct and the barcode RNA leads the RNA to covalently link to the fusion protein. **B**. Cells expressing 6X-His-TrmA (E358Q) or 6x-His-FLAG-TrmA (E358Q) were pooled at equal optical density, and lysates were bound to Ni-NTA or to anti-FLAG M2 antibody, followed by barcode sequencing. Stacked bar plots show the proportions of the input barcode read count based on fusions that enrich on Ni-NTA beads, and the output barcode read count based on fusions that enrich on anti-FLAG M2 beads. The average ratio of 6X-His-FLAG-TrmA (E358Q) to 6X-His-TrmA (E358Q) was ∼2.5 in the input (Ni-NTA) and increased to ∼110 after selection on anti-FLAG M2 beads (prop test: p-value <10^−2^ □, n=2), corresponding to an average ∼43-fold enrichment (ratio of output [6X-His-FLAG-TrmA (E358Q) / 6X-His-TrmA (E358Q)] g for high throughput assessment of the effects of all sing **C**. TrmA (E358Q) fusion proteins were expressed in mammalian cells. The ratio of output [WT-FLAG-TrmA (E358Q)-TPMT/Mutant-FLAG-TrmA (E358Q)-TPMT]/input [WT-FLAG-TrmA (E358Q)-TPMT/Mutant -FLAG-TrmA (E358Q)-TPMT] was calculated. The stacked bar plot shows the proportions of barcode read counts from plasmid DNA isolated from mammalian cells (input) and from fusions that enrich on anti-FLAG M2 beads (output) (prop test: p-value 10^−2^ □, n=3).

To demonstrate that addition of the FLAG tag, an epitope of eight amino acids that binds to the M2 monoclonal antibody^18^, enabled efficient recovery of TrmA (E358Q) fusions, we generated two constructs. One encoded 6X-His-TrmA (E358Q) and the second encoded 6x-His-FLAG-TrmA (E358Q), each with a unique barcode sequence. These constructs were expressed individually in *E. coli*, and the lysates were bound to Ni-NTA beads (selecting for binding of the 6X-His tag), followed by reverse transcription of the linked barcode and qPCR analysis. We found that both the 6X-His-TrmA (E358Q) and 6X-His-FLAG-TrmA (E358Q) bound to the Ni-NTA beads (Fig. S2). However, by using beads coupled to the anti-FLAG M2 antibody (selecting for binding of the FLAG epitope), we observed that only the FLAG-tagged construct was efficiently bound (with a 16-fold enrichment) (Fig. S2). Using DNA sequencing, we found that the FLAG-tagged construct was enriched ∼42-fold compared to the construct without this tag (Fig. 1B).

Two additional features of the assay were tested. First, to assess whether non-cognate barcodes liberated from neighboring cells during cell lysis link to free TrmA (E358Q) fusion proteins, we tested the addition of 10 µM stem-loop sequence (with no barcode) during lysis. The stem-loop addition cut almost in half the small amount (1.4%) of either linking of non-cognate barcodes or nonspecific binding to the NTA beads of the fusion protein without a tag (Fig. S3 and Supplementary discussion). Second, we tested whether a 6X-His version of an engineered TrmA bound more specifically to an engineered stem-loop^19^ than to the WT stem-loop, and found that it did not (Fig. S4 and Supplementary discussion). Based on these results, in the experiments below, we added 10 µM stem-loop without a barcode during lysis and used the TrmA (E358Q) mutant.

We sought to determine whether Dosa works in mammalian cells, as *E. coli* lacks eukaryotic-specific chaperones that may be critical for folding of some proteins. To assess whether stem-loop RNA barcodes can link to TrmA (E358Q) and be specifically enriched in this context (Fig. S5A), mammalian cells were transfected with plasmids encoding either the RNA barcode alone, the FLAG-tagged human thiopurine methyltransferase (TPMT) fused to TrmA (E358Q), or both. TPMT was chosen because of an available dataset from deep mutational scanning^4^. To test whether TrmA (E358Q) tolerates fusion at different positions, TPMT was fused to either the N- or C-terminus of TrmA (E358Q) (Fig. S5B). Enrichment on anti-FLAG M2 beads followed by qPCR showed that when the stem-loop RNA barcode alone was expressed without the TrmA (E358Q) fusion protein, the RNA did not enrich on anti-FLAG M2 beads (Fig. S5C). Enrichment of the RNA barcode when co-expressed with FLAG-TrmA (E358Q) fused to TPMT in either orientation was about ∼1,000 fold (ΔCq ∼10 cycles, n=2) compared to RNA expressed alone. This result indicates that a stem-loop barcode was found on the antibody beads only when TrmA (E358Q) was also present in the reaction.

We tested whether barcoded proteins expressed in mammalian cells could be specifically enriched. Two fusion constructs with unique barcode sequences were generated, one with WT FLAG-TrmA (E358Q)-TPMT and an identical one except that the FLAG tag contained mutations that prevent its binding to M2 antibody and therefore enrichment on beads. Transfected cells were pooled, and lysates were enriched on anti-FLAG M2 beads. Barcode sequencing showed that the ratio of WT-FLAG to FLAG mutant was enriched ∼6.8-fold (Fig. 1C). Together, these results demonstrate that the covalent linking of an RNA barcode to a protein required for the Dosa approach is amenable to mammalian cells.

### The Dosa approach recapitulates protein-protein interaction binding data

The FLAG epitope (DYKDDDDK) is widely used for Western blots, protein purification and ELISA assays^18^ and its binding to the anti-FLAG M2 antibody serves as a model protein-protein interaction. We used Dosa to assess the effect of mutations in the FLAG peptide on antibody binding. These experiments were motivated by the availability of a co-crystal structure of the FLAG peptide bound to the anti-FLAG M2 antibody^20^ and a dataset from another high throughput assay, CasPlay, of FLAG peptide variants^14^.

A site saturation library of 152 FLAG epitope variants was generated using an NNK-codon oligonucleotide pool that was cloned between an N-terminal 6X-His tag and TrmA (E358Q). The median number of barcodes per mutant was 10, with a total of ∼1,900 unique barcodes. The invariant N-terminal 6X-His tag of the library construct allowed the capture of these fusion proteins on Ni-NTA beads, providing the abundance of each of the protein variants in the pool (the input for enrichment score calculation). However, only FLAG epitope variants capable of binding to the anti-FLAG M2 antibody were enriched using magnetic beads coupled to the antibody (the output for the enrichment score) (Fig. 2A). A log_2_ ratio of [barcodes bound to Ni-NTA beads (Input) / barcodes bound to anti-FLAG M2 beads (output)] provides the enrichment score.

**Fig. 2.**
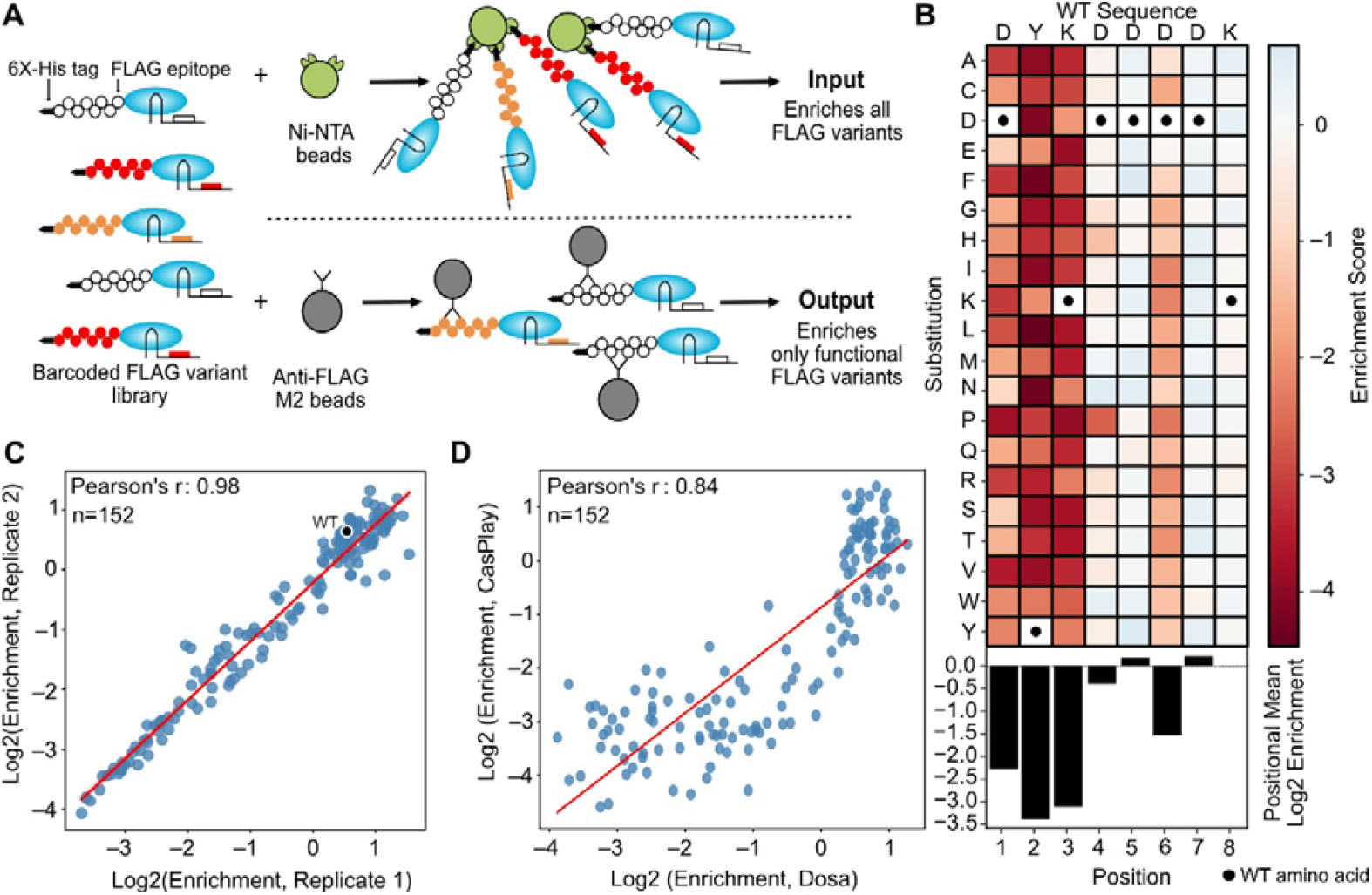
Mutational analysis of the FLAG epitope for its binding to anti-FLAG M2 antibody. **A**. Schematic representation of the experimental workflow. Lysates with TrmA (E358Q)-barcoded proteins are bound to Ni-NTA beads, and the bound barcodes are reverse transcribed and amplified for sequence analysis; all barcoded proteins contain a 6X His-tag and enrich on Ni-NTA beads (input). On anti-FLAG M2 antibody beads, only functional FLAG variants bind, constituting the output. **B**. Heatmap illustrating the effect on FLAG-antibody binding of amino acid substitutions at each position within the FLAG epitope; black dots are wild type residues. The bar graph shows the positional mean of all amino acid substitutions at each position. **C**. Pearson’s correlation plot between two biological replicates; each dot is a unique variant. Red line indicates linear regression fit. **D**. Pearson’s correlation plot comparing the FLAG epitope mutational scan data from the Dosa method to enrichment values obtained using the CasPlay method^14^; dots and red line as in C.

A heatmap describing how single variants in the FLAG peptide affect binding to the anti-FLAG M2 antibody (Fig. 2B) allowed us to compare data from Dosa to the CasPlay^14^ method, which analyzed FLAG epitope binding to the same antibody using dCas9 fusions. The mutational data from Dosa show both good reproducibility (Pearson’s r = 0.98) (Fig. 2C) and agreement with the CasPlay mutational scan (Pearson’s r = 0.84) (Fig. 2D). The shape of the curve (Fig. 2D) indicates that Dosa is less sensitive than CasPlay at discriminating variants with near-wild type values, but Dosa shows a much wider range of scores for FLAG variants with poor binding.

The Dosa data indicate that there are strong limitations on which residues can be varied without affecting anti-FLAG antibody binding. The heatmap shows that positions D1, Y2, K3 and, to a more limited extent, D6, were intolerant of substitutions (positional enrichment means of −2.2, −3.3, −3.1 and −1.5, respectively). The crystal structure of the FLAG peptide bound to the anti-FLAG M2 antibody^20^ shows that positions D1, Y2, K3, D4 and D6 are involved in binding to the antibody, consistent with the enrichment scores from Dosa. Although D4 forms a double salt bridge with R32 in the antibody light chain, the Dosa mutational data, in agreement with the results from CasPlay,^14^ indicate that most substitutions at this position were tolerated. R32 in the light chain undergoes conformational change upon antigen binding and may be able to accommodate mutations, apart from proline, at D4^20^. The side chain of D5 points away from the antibody, indicating that it is likely not critical for binding, as seen in the mutational data. The lack of electron density of D7 and K8 in the crystal structure and the observation that truncation of the peptide that eliminates positions 7 and 8 leads to only modest reduction in affinity^20^ are consistent with these positions being dispensable for interaction.

### Measurement of protease substrate specificity using Dosa

We next applied Dosa to determine the substrate specificity of a protease. Enteropeptidase, also known as enterokinase, is an intestinal enzyme that activates trypsinogen to trypsin^21,22^. Trypsin activates other zymogens, such as chymotrypsinogen, proelastases and prolipases, for their roles in protein digestion. Enteropeptidase cleaves trypsinogen at its activation peptide, whose sequence is DDDDK^22^ (Fig. S6A). Three variants (shown in Fig. S6A) in the trypsinogen activation peptide have been characterized biochemically for their activation by enteropeptidase:^23,24^ D19A is moderately more active than WT, D22G is severely compromised for cleavage and K23R is considerably more active for cleavage than the WT.

Because the C-terminal five residues of the FLAG peptide (DDDDK) are identical to those in the activation peptide of enteropeptidase, we were able to use the same FLAG peptide library of 152 variants used to assay antibody binding to profile enteropeptidase substrate specificity (Fig. 3A). The library of barcoded substrates was produced in *E. coli*, purified, immobilized on Ni-NTA beads and treated with enteropeptidase. Cleavage of the substrate led to loss of its corresponding barcode from the beads during washing, whereas a variant substrate that resists cleavage resulted in its barcode being retained on beads (Fig. 3A). Thus, barcodes corresponding to the cleavable substrates were depleted, and barcodes corresponding to uncleavable substrates were enriched. We computed the scores such that variants that cleave as well or better than WT have a positive enrichment score, and those resistant to cleavage have a negative enrichment score.

**Fig. 3.**
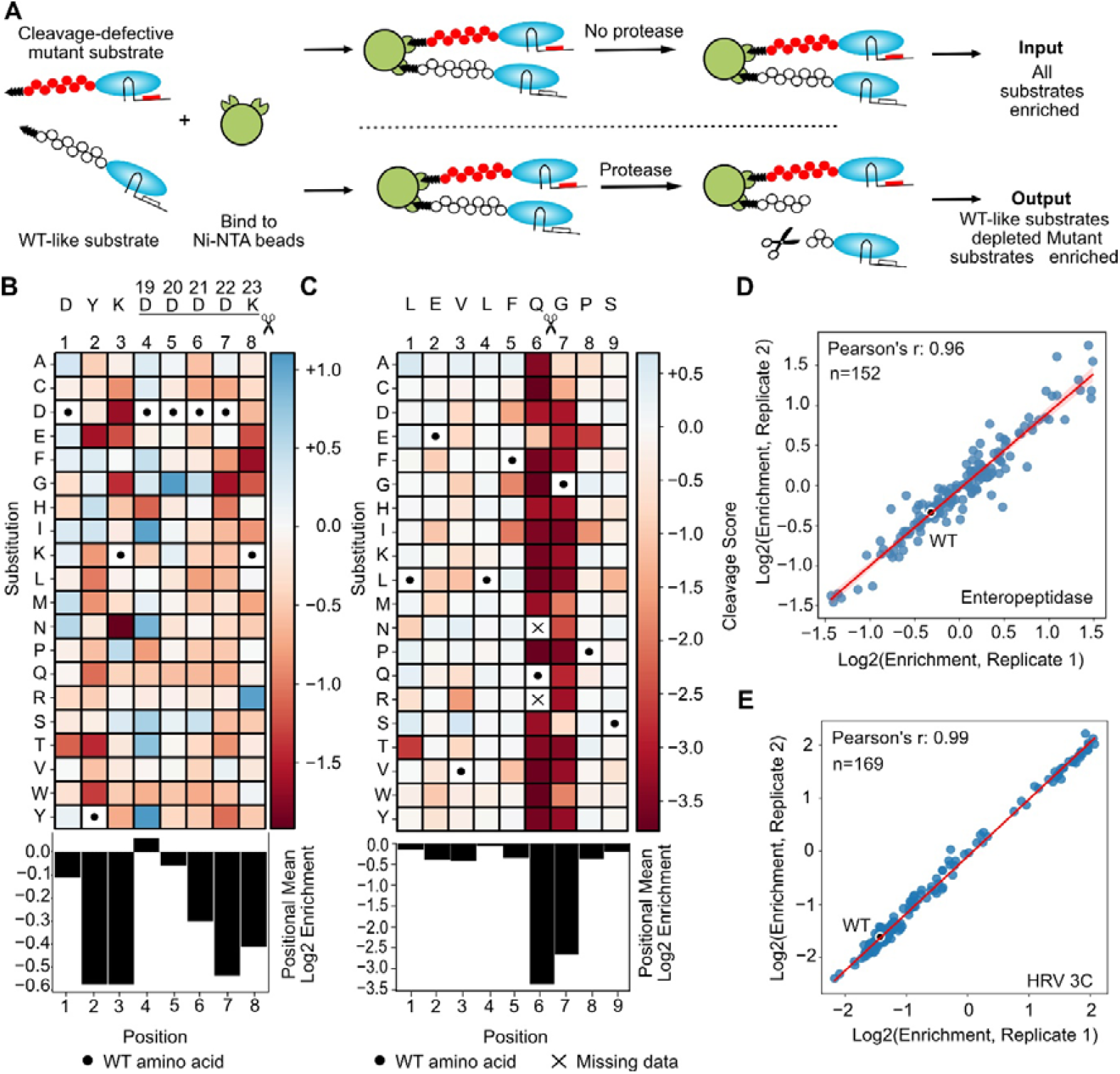
Assay for protease cleavage site preferences. **A**. Schematic representation of the assay. Barcoded proteins are immobilized on Ni-NTA beads, and a recombinant protease is added. Following incubation, wild-type and mutant proteins with enhanced cleavage are depleted from the beads during washing steps, while variants that do not undergo cleavage are retained on the beads. **B**. Heatmap showing FLAG epitope cleavage by enteropeptidase. Scissors indicate the cleavage site. The underlined sequence corresponds to the activation peptide in trypsinogen and the numbers above it (19-23) correspond to positions in trypsinogen. The bar plot below the heatmap displays the positional mean of substitutions at each position. **C**. Heatmap showing cleavage of a substrate by the human rhinovirus 3C protease. Scissors indicate the cleavage site. The bar plot displays the positional mean of substitutions. **D**. Pearson’s correlation plot between two biological replicates of the enteropeptidase cleavage assay. Each dot is a unique variant. Red line indicates linear regression fit. **E**. Pearson’s correlation plot between two replicates of the rhinovirus 3C protease cleavage assay; dots and red line as in D.

Substitutions in D1, D4, and D5 of the DYKDDDDK FLAG peptide had relatively minor effects on cleavage, with positional means of −0.11, +0.11 and −0.05, respectively (Fig. 3B). D6, D7 and K8 were intolerant to substitutions, consistent with previous *in vitro* experiments^25^. We compared the Dosa dataset to the small set of variants found in clinical conditions^23^, including K23R (in trypsinogen numbering, Fig. 3B) implicated in hereditary pancreatitis, that have been analyzed biochemically^23,24^ (Fig. S6B). By the Dosa assay, D19A cleaved slightly better (score +0.58) than the WT sequence (+0.32), D22G cleaved very poorly (−1.26) and K23R cleaved much better than WT (+1.36), yielding a Pearson’s correlation between Dosa scores and biochemical analyses of r = 0.96 (Fig. S6B). The glycine substitution at D4, D5, and D6 in the DYKDDDDK peptide also performed better than WT (Fig. 3B), likely due to its making the peptide more flexible and consistent with a random peptide screen of enteropeptidase substrates^26^ that enriched for glycine at D6.

We assayed the effects of substitutions in another protease cleavage site. Human rhinovirus (HRV) belongs to a family of medically important viruses known as the picornaviridae, which also include poliovirus and hepatitis A virus. This family of viruses produces a protease known as 3C, which acts in viral maturation by cleaving the virus-encoded polyprotein^27^. In addition, the 3C protease cleaves host proteins to modulate host responses, including translation^28^ and immunity^29^.

We generated a library of 178 single amino acid substitutions (two variants were not generated) in the optimal substrate peptide LEVLFQGPS^30^ for human rhinovirus 3C protease with an N-terminal 6X-His tag and C-terminal TrmA (E358Q) protein. The Q6 and G7 positions were intolerant to all substitutions, with positional means of −3.3 and −2.6, respectively (Fig 3C), as expected given that cleavage occurs between these two amino acids. A modified yeast surface display method, yeast endoplasmic reticulum sequestration screening (YESS), was used to profile substrate specificities at the Q6, G7, and P8 positions^30^. We found good correlation between the 3C protease substrate specificity at these three positions found by Dosa and by the YESS approach^30^ (Pearson’s correlation r = 0.87) (Fig. S7A).

An alignment of 3C cleavage sites in the HRV polyprotein indicates that position G7 tolerates serine (Fig. S7B), consistent with the Dosa mutational data (Fig 3C).The YESS and Dosa approaches also showed that serine, cysteine, and alanine substitutions at G7 were cleaved, albeit with lower efficiency^30^. The Dosa mutational scan did not identify any 3C variants that were cleaved more than marginally better than the WT substrate, confirming that the peptide substrate used in the scan was highly optimized^30^. Biological replicates of the enteropeptidase cleavage assay (Pearson’s r = 0.96) (Fig. 3D) and 3C assay (Pearson’s r =0.99) (Fig. 3E) correlated well. The Dosa results for cleavage specificity of both enteropeptidase and human rhinovirus 3C protease were highly correlated with the limited data available from previous studies, validating Dosa for determining protease cleavage sites.

### Dosa assay to assess protein solubility

We leveraged the covalent linkage between a protein and its RNA barcode in Dosa to develop a massively parallel protein solubility assay. The solubility of a protein is often assessed by isolating soluble and insoluble fractions via centrifugation, with each of these fractions then separated by SDS-PAGE to quantify the relative band intensity of the protein; soluble proteins preferentially enrich in the supernatant, whereas insoluble ones enrich in the pellet. Proteins in the insoluble fraction can be solubilized, typically by denaturing conditions using SDS or chemical denaturants such as urea. Proteins barcoded in the Dosa method retain their RNA barcode even after solubilization in 8M urea, and 6X-His proteins can be enriched on Ni-NTA beads even under denaturing conditions.

To benchmark the assay, we used *E. coli* maltose binding protein (MBP), a globular protein that has been studied extensively, including the assessment of mutants that alter solubility^31^. G19C has WT-like solubility, whereas Y283D and the double mutated G32D, I33P are insoluble^32^. Using a fusion protein of TrmA (E358Q) with WT MBP; with the G19C and Y283D single mutants; and with the double mutant G32D, I33P, a gel-based solubility assay showed that G19C was comparably soluble as WT and that the other two mutants were insoluble (Fig. S8); however, given the presence of multiple endogenous bands on the gel, it was not possible to accurately quantify solubility by this assay. The similarity of the gel results to the previous solubility assay^31^ indicates that the TrmA fusions did not have any major effects on the solubility of these MBP variants.

Having established feasibility, we used Dosa to determine the solubility of these same MBP variants. *E. coli* cells expressing a pool of WT and the three MBP mutants fused to TrmA (E358Q), each expressing a unique barcode, were lysed. After separation of the soluble proteins, the insoluble fraction was resuspended in 8M urea to solubilize the proteins. MBP proteins isolated from both fractions were enriched on Ni-NTA beads, and barcodes were reverse transcribed and sequenced. The log_2_ ratio of the barcodes present in the soluble and insoluble fraction provides a score of protein solubility. The Dosa results (Fig S9) indicate that the solubility results were largely concordant with those from the gel-based assay.

Aβ(1-42), a peptide associated with Alzheimer’s disease, is one of the cleavage products of amyloid precursor protein through the sequential action of β-secretase and γ-secretase^33^. Aβ(1-42) is aggregation-prone and forms amyloid fibrils found abundantly in plaques of brains isolated from Alzheimer’s disease patients. A double mutant [Aβ(1-42), F19S, L34P], called GM6, does not aggregate and largely remains in the soluble fraction^34^. 6X-His Aβ(1-42) was expressed as a fusion protein at the N-terminus of TrmA (E358Q), and barcoded proteins were expressed in *E. coli*. The gel-based solubility assay with WT Aβ(1-42) and the GM6 Aβ(1-42) fused to TrmA (E358Q) showed that GM6 was more soluble than WT (Fig. S10A). We also validated that the GM6 double mutant was more soluble than WT using the Dosa assay (Fig. S10B).

We sought to use Dosa to measure the solubility of a variant library of Aβ(1-42) (Fig. 4A). Single amino acid substitutions were made in Aβ(1-42) fused to TrmA(E358Q), and a solubility score – as for MBP – was calculated for 794 variants (Fig. 4B), with a replicate correlation Pearson’s r = 0.81 (Fig. S11A). Comparison to a published solubility screen of Aβ(1-42) carried out in yeast showed a correlation of Pearson’s r = 0.59 (Fig. S11B). Differences between the assay results could be caused by different expression systems and different fusion proteins, as Dosa measured solubility whereas the other assay employed a yeast DHFR aggregation assay that uses growth as a proxy for aggregation^3^.

**Fig. 4.**
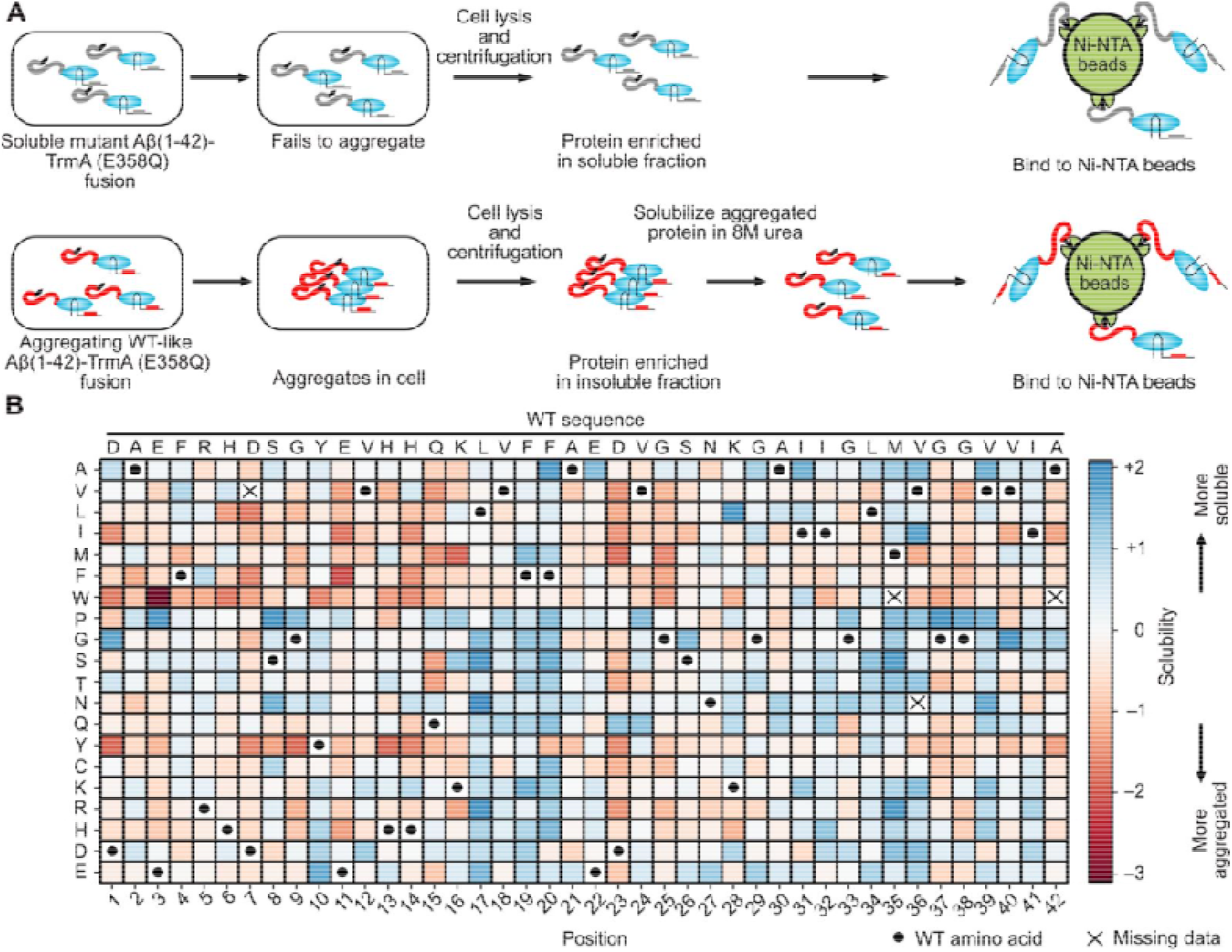
Assay of protein solubility. **A**. 6X-His Aβ(1-42) mutants fused to TrmA (E358Q) are expressed in *E. coli*. The WT Aβ(1-42) has a propensity to aggregate whereas variants that do not aggregate are more soluble. Cell lysis and centrifugation separate soluble and insoluble fractions, and the insoluble fraction containing aggregates is solubilized in 8M urea. The barcoded proteins from each fraction are enriched on Ni-NTA beads, and the barcodes are sequenced. **B**. Heat map showing the effect of substitutions at each position on Aβ(1-42) solubility.

The positional means of the solubility scores indicate that, on average, substitutions at positions 17-20 and 34-36 resulted in more solubility, and substitutions at positions 7, 11, 23, 25, 37 and 38 led to more aggregation (Fig. S12A). Substitution with hydrophobic amino acids led to more aggregation than substitution with hydrophilic ones (Fig. S12B, C). Consistent with earlier observations^3^, proline substitution resulted in the highest solubility score (median = 0.49). Tryptophan, tyrosine, and phenylalanine substitution had the opposite effect, resulting in increased aggregation, with medians of −0.7, −0.68 and −0.53, respectively (Fig. S13).

The cryo-EM structure of amyloid Aβ(1-42) isolated from patients with Alzheimer’s disease has found two types of filaments, type I from patients with sporadic Alzheimer’s disease and type II from those with the familial form.^35^ Positions 17-20 and 34-36 cluster together structurally to forms a hydrophobic patch in type I filaments. E22 and D23 coordinate an unknown metal ion (in both types of filaments). Substitutions at D23 tend to increase aggregation (Fig. 4B). The type II filament interface is stabilized by the amino group of K28 from one monomer to the backbone carboxyl group of A42 from a second monomer. Polar substitutions at K28 increase solubility (Fig. S12C). Our mutational data support the formation of type I filaments in *E. coli*, although only structural analysis can confirm this finding. Of 11 familial variants in amyloid Aβ(1-42)^36^, the Dosa dataset includes all but the E22Δ deletion. Seven of the ten variants were more insoluble than the WT (H6R, D7H, A21G, E22G, E22Q, D23N, L34V); two of the other three showed no change (A2V, K16N) and one showed increased solubility (E22K) (Fig. 4B). Solubility in the Dosa assay is only one measure of Aβ(1-42) variant properties, which show complex aggregation kinetics and multiple morphological conformations and immunological reactivities in other assays^36^.

## Discussion

Here, we introduce a protein display method in which RNA barcodes are covalently linked to proteins *in vivo* via the *E. coli* TrmA (E358Q) mutant. This covalent RNA-protein link enables proteins to be subjected to biochemical assays – including those, like the solubility assay used here, that require harsh conditions such as high temperature or a strong denaturant. Moreover, the method can be applied to proteins that require a cellular environment for proper folding or post-translational modification. The barcoding approach works both in *E. coli* and human cells, indicating that it is likely to be usable with most transformable organisms. TrmA (E358Q) is highly specific for linking to its stem-loop sequence, and at 42 kDa, it is much smaller than Cas9 (158 kDa) used in a related approach that results in noncovalent binding of an RNA to a hybrid protein^14^. Another useful feature are the multiple barcodes associated with each variant, indicating the degree of variability in measurements.

One limitation of Dosa is that proteins subjected to analysis are not native; instead they are hybrid proteins that include both a TrmA (E358Q) domain and an RNA. While less likely to impact a protein interaction site or cleavage site, the TrmA (E358Q) and RNA components of a hybrid protein could affect the folding or solubility of the protein of interest. However, numerous studies that use green fluorescent protein and other reporter proteins indicate that reporter activity is well correlated with the folding and solubility of the upstream protein of interest^37^.

The barcoding RNAs in the current version of Dosa are separately encoded from the mRNAs, preventing facile recovery of the mRNA of a protein with sought-after properties. This separate encoding also does not allow for successive rounds of protein selection as can be done with methods such as phage display. However, it should be possible to tag whole mRNAs by incorporating the stem-loop into the 3’ untranslated region of the mRNAs, enabling both biochemical selection and the recovery of mRNAs for subsequent rounds of cloning and selection. Dosa could also be expanded to barcode the entire proteome of an organism and subject it to assays. Other improvements could include engineering TrmA (E358Q) to covalently link *in vivo* to a completely orthogonal stem-loop sequence and reducing the size of TrmA (E358Q) without compromising its catalytic activity or stability, to minimize its effect on the protein of interest. The use of a more thermostable TrmA should allow higher assay temperatures to be employed. Finally, the large datasets of activities for proteins and protein variants resulting from this method should enable the improvement of machine learning methods that assess and predict biochemical activities.

## Materials and Methods

### Cloning

For TrmA protein purifications, N-terminal 6X-His tagged TrmA WT and mutants (E358Q) and (E358Q, C234A) were cloned into pNIC28-BSA4 (Addgene # 26103) using Gibson assembly cloning. During the initial stages of Dosa method development in *E. coli*, we used pNIC28-BSA4 vector (Fig. 1 and related supplements). The sequence of the 19-base stem-loop 5’ GCTGTGTTCGATCCACAGC 3’ was expressed separately from a T7 promoter with flanking primer sequences for amplification (Forward 5’ ATAATCATGAGTGCCGCG 3’ and Reverse 5’ AACTGGACGGAACTCGAGA 3’). A random 10-base barcode was inserted between the primer amplification sites.

pCDNA 3.1 (Thermo) was the vector for mammalian expression. Gibson assembly cloning was performed to generate fusion constructs with the TrmA (E358Q) and human thiopurine methyl transferase. A linker sequence ‘ENLYFQSGSG’ was cloned between the fusion proteins. The RNA barcode was expressed from a human U6 promoter that generates a transcript with the 19 base tRNA stem-loop TrmA recognition site, a 10-base unique barcode and PCR priming sites (as described above).

#### Library cloning

Detailed vector maps and primer sequences are in supplemental data. Library experiments were performed using pET-2A-T (Addgene #29665), which contains an ampicillin resistance marker. This vector backbone was used after removal of Esp3I and BsaI restriction sites. 6X-His TrmA (E358Q) was expressed from a T7 promoter. The sequence between the 6X-His tag-TEV recognition site and TrmA (E358Q) contained a stuffer fragment that contained two BsaI sites for variant library insertion. The RNA barcode was transcribed from a T7 promoter with the 19-base stem-loop and primer sequences as described above. Another DNA stuffer fragment with two Esp3I sites was inserted between the priming sites for the RNA barcode that allowed barcode ligation in subsequent steps. Library cloning was performed in two steps, the first inserting barcodes and the second inserting the genes encoding protein variants (Fig. S14).

1. <u>Generation of pre-barcoded plasmid library</u>. pET-2A-T (described above) was used. An Ultramer sequence 5’ TATGCGAATTCCTGCTATCGCGTCTCACTATGTGTCTAGCTATTAGTAATACGACTCACTATAGGGGGCTGTGTTCGATCCACAGCATAATCATGAGTGCCGCG 3’ and a primer 5’ TAATCATGAGTGCCGCGNNNNNNNNNNTCTCTGAGACGGCTAGTTAC 3’ (N represents an equal mix of all four nucleotides) was purchased from IDT. They were annealed by heating to 90°C and cooled. Klenow DNA polymerase (NEB) was used to fill in the single-stranded DNA. The products were purified on Zymo-Spin I column (Zymo Research) and ligated to the vector using Esp3I sites. A library of ∼5 million barcoded plasmids was generated. After ligation of barcodes to the vector, Esp3I sites are no longer present in the plasmid. This pre-barcoded library was used to generate mutational scanning libraries.
2. <u>Cloning of variant protein library.</u> The variant library DNA lacked any BsaI or Esp3I sites. The DNA included a 5’ adaptor (5’ TTCGCACTGTCAGGGTAGGTCTCATGGGT 3’) and 3’ adapter (5’ ACGCTAGTGCAAAACACCGGTCTCGTTCTC 3’) that contained BsaI sites for Golden Gate cloning and primer binding sites for amplification. Oligonucleotides encoding variants of the FLAG epitope and the HRV 3C substrate were purchased as NNK [where N is a mix of all four nucleotides, and K is G or T] oligonucleotide pools from IDT and made double-stranded using a single cycle of Kapa Hot start polymerase using the primer 5’ ACGCTAGTGCAAAACACCG 3’. Aβ(1-42) oligonucleotide pools (purchased from Twist Biosciences) were amplified by Kapa Hot start polymerase using primers 5’ TTCGCACTGTCAGGGTAG 3’ and 5’ ACGCTAGTGCAAAACACCG 3’. Amplification was monitored by qPCR and the PCR was stopped 3 cycles after Cq. The reactions were purified over a Zymo-Spin I column. The inserts were ligated into the BsaI site of the pre-barcoded vector (see step 1). Following another Zymo-Spin I column cleanup, the DNA was transformed into *E. coli* ElectroMAX™ DH10B cells (Thermo) to generate libraries. Libraries were bottlenecked to typically 10-20 times the expected number of variants to generate 10-20 barcodes per variant. Subassembly of plasmids was performed using PacBio long read sequencing, after linearization of 3C substrates and Aβ(1-42) libraries with I-SceI and the FLAG epitope library with BamHI. Linearized plasmids were processed using SMRTbell® Express Template Prep kit 2.0 as described by the manufacturer and barcoded with different adapters (Barcoded overhang adapter kit 8A, PacBio).

To generate constructs used in model selections with MBP (WT, G19C, Y283D, G32D/I33P), PCR fragments were ligated to the pre-barcoded plasmid library (Step 1 of library cloning). The WT and GM6 [F19S/L34P] Aβ(1-42) were synthesized as Ultramer DNA (IDT) and cloned as described for MBP. Resulting constructs of MBP and Aβ(1-42) had an N-terminal 6X-His tag.

### TrmA protein expression and purification

N-terminal 6X-His tagged TrmA WT and mutants (E358Q) and (E358Q, C234A) were cloned in pNIC28-BSA4. BL21(DE3)-transformed plasmids were grown with 50 µg/ml kanamycin overnight in LB broth. The next day, fresh cultures (25 ml) were inoculated with 0.1 O.D of the overnight culture in LB kanamycin. Upon the cultures reaching O.D of 0.6, IPTG was added to a final concentration of 0.1 mM. Cultures were grown for 2 more hr at 30°C and then cell pellets were harvested by centrifugation and frozen at -20°C until purification. To the frozen cells, Xtractor buffer (#635671, Takara) and protease inhibitor 0.1 mM Pefabloc (#PEFBSC-RO, Roche) were added, followed by incubation at room temperature for 15-20 min. Cells were centrifuged at 16,000 rpm for 30 min. Supernatants were bound to pre-equilibrated His-Pur Ni-NTA agarose (# 88221, Thermo) for 60 min. The flow-through was discarded and the beads were washed with wash buffer I (50 mM Hepes pH 7.5, 0.5 M NaCl, 10% glycerol, 0.5 mM TCEP, 10 mM imidazole) and then wash buffer II (50 mM Hepes pH 7.5, 0.5 M NaCl, 10% glycerol, 0.5 mM TCEP, 25 mM imidazole). Proteins were eluted in elution buffer (50 mM Hepes pH 7.5, 0.5 M NaCl, 10% glycerol, 0.5 mM TCEP, 350 mM imidazole). Eluted samples were run on SDS-PAGE to verify purity. Samples were dialyzed overnight in storage buffer (20mM Hepes pH 7.4, 50mM NaCl, 0.5 mM TCEP) and flash frozen and stored at −70°C or run in SDS-PAGE, as needed.

### Western blotting

Samples were fractionated by SDS-PAGE and transferred onto nitrocellulose membranes (#1704158, Bio-Rad). Membranes were blocked with 1% BSA (#A-420-50, Gold Bio) in Tris buffered saline pH 7.4, 0.05% Tween 20 (TBST) and incubated at room temperature for 1 hr with shaking. Blocked membranes were incubated with anti-6X-His antibody (MA-135-A488, Thermo) for 1 hr at room temperature. Membranes were washed with TBST three times each for 10 min before imaging using ChemiDoc™ MP imaging system (Bio-Rad).

### Anti-FLAG M2 antibody binding assay

#### Production of barcoded proteins

Electrocompetent *E. coli* BL21 (DE3) cells were transformed with the plasmid library. Cells were recovered and grown overnight in LB with 100 µg/ml ampicillin. The next day the cells were diluted to an O.D of 0.1 and grown at 37°C until the O.D reached 0.5. IPTG was added to a concentration of 0.1 mM and cells were grown further for 2 hr at 30°C. Cell density was quantified and aliquots of culture corresponding to a total of 2 O.D were collected and flash frozen after removal of media.

#### Output enrichment (binding to anti-FLAG M2 beads)

150 ml Xtractor buffer (Takara) was added to 2.0 O.D of frozen cells. 0.1 mM Pefabloc, 10 µM 19 base stem-loop, and 1 µl murine RNase inhibitor (NEB) were added, and the cells were thoroughly resuspended and then incubated on ice for 20 min. The supernatant (clarified lysate) was obtained by centrifugation at 15,000 rpm for 30 min at 4°C. 50 µl of M2 anti-FLAG beads (M8823, Sigma) were equilibrated with wash buffer (50 mM Tris pH 7.5, 300 mM NaCl). The beads were added to the supernatant and incubated with rotation in the cold room for 40 min. Using a magnetic rack, beads were separated from lysate and washed thoroughly with 2X 500 µl wash buffer, followed by one wash of 250 µl low salt buffer (50mM Tris pH 7.5, 50 mM NaCl).

#### Input enrichment (binding to Ni-NTA beads)

A clarified lysate was prepared as in output enrichment and bound to His-Pur™ Ni-NTA beads (Thermo) using 30 µl (wet beads) beads equilibrated with imidazole wash buffer (50 mM Tris pH 7.5, 300 mM NaCl, 25 mM imidazole) and incubated with rotation in the cold room for 40 min. The beads were separated from lysates using centrifugation at 4,000 rpm for 1.5 min The beads were washed twice with 500 µl imidazole wash buffer, once with 250 µl wash buffer (see output enrichment for composition) and once with 250 µl low salt buffer.

### Reverse transcription, sample indexing and short read sequencing

Barcoded proteins bound to beads (both input and output from the FLAG epitope binding assay) were used. The beads were resuspended in 40 µl water and 5 µl 10X DNase I buffer (Thermo) along with 2 µl DNase I (Thermo) and 1 µl RNase inhibitor (NEB) and incubated at 37°C for 1 hr. DNase I was inactivated by adding 0.6 mM EDTA and heating the reaction to 75°C for 10 min. 1 µl additional RNase inhibitor was added.

3 µl of the sample was used as input for reverse transcription by SuperScript IV (Thermo) using the manufacturer’s guidelines. The oligonucleotide for priming the reverse transcription was 5’ ACGAAAACTGGACGGAACTCGAG 3’. 1 µl RNase H was added and incubated at 37°C for 20 min. cDNA was used as template for amplification of the barcodes and addition of P7 and P5 adaptors and index sequences for Illumina short read sequencing (see Supplementary data for primers). Amplification was monitored by using 1X SYBR Green I (S7563, Thermo) in the reaction. SYBR Green I was diluted into a working stock of 100X in DMSO and diluted 10X in water immediately prior to preparing qPCR reactions. Typically, 10-20 cycles were used for amplification. Samples without reverse transcription were used as control for monitoring qPCR. Products were purified on Zymo columns and quantified with Qubit (Thermo). Library quality was assessed using TapeStation (Agilent) with an expected size of the DNA of 161 bp. The setup of the sequence used for sequencing and the annealing sites for the primers is shown in Supplementary data (genbank file). Libraries were sequenced on a Nextseq2000 (Illumina). Custom primers were used for sequencing (see Supplementary data for custom primer sequences).

### Protease substrate specificity assay

#### Production of barcoded proteins

Barcoded proteins were produced in *E. coli* as described for the anti-FLAG M2 antibody binding. The 3C substrate library was produced identically except that 1 mM IPTG was used for protein induction.

#### Enteropeptidase cleavage assay (output)

Barcoded proteins were isolated on Ni-NTA beads as described for the anti-FLAG M2 antibody binding. After 2X washing with imidazole buffer, the beads were washed with enterokinase cleavage buffer (20 mM Tris pH 7.5, 50 mM NaCl, 2 mM CaCl_2_). 5 µl enteropeptidase (Enterokinase [Bovine], #P8070S NEB) was added and incubated at 4°C overnight with 2 µl murine RNase inhibitor (NEB). The beads were washed twice with 500 µl enterokinase cleavage buffer (20 mM Tris pH 7.5, 50 mM NaCl, 2 mM CaCl_2_), followed by one wash with 250 µl 50mM Tris pH 7.5, 50 mM NaCl. Beads were processed as described in Reverse transcription, sample indexing and short read sequencing.

#### 3C cleavage assay (output)

Barcoded proteins were isolated on Ni-NTA beads as described for anti-FLAG M2 antibody binding. After 2X washing with imidazole buffer, the beads were washed with 3C protease cleavage buffer (50 mM Tris pH 7.5, 150 mM NaCl, 0.5 mM TCEP). 5 µl recombinant HRV 3C protease (2U/µl) (# 27084301, Cytiva) was added along with 2 µl murine RNase inhibitor and incubated with rotation for ∼16 hr at 4°C. The beads were washed twice with 500 µl cleavage buffer, followed by one wash with 250 µl 50mM Tris pH 7.5, 50 mM NaCl. The beads were then processed as described for Reverse transcription, sample indexing and short read sequencing.

#### Input sample (both proteases)

The identical protocol described for the output sample was followed without the addition of the recombinant protease.

### Protein solubility assay

#### Production of barcoded proteins

Aβ(1-42) barcoded proteins were produced with the same protocol as the FLAG epitope barcoded proteins. MBP constructs were induced either with 0.1 or 1 mM IPTG at 37°C for 2 hours.

#### SDS-PAGE solubility assay

The O.D. of induced cultures [either 1 mM or 0.1 mM IPTG for MBP-TrmA (E348Q) fusions and 0.1 mM for Aβ(1-42)-TrmA (E358Q)] was quantified and cells corresponding to 4 O.D of culture was pelleted and flash frozen. 200 µl Xtractor buffer was added and cells were lysed on ice for 20 min with 0.1 mM Pefabloc. The supernatant-containing soluble fraction was obtained by centrifugation at 21,000 g for 30 min. The pellet was resuspended in an equal volume of Xtractor buffer. Equal volumes of samples from the supernatant and pellet fractions were loaded onto SDS-PAGE gels.

#### Dosa solubility assay

Cells were lysed and the soluble proteins present in the clarified supernatant were isolated. To the pellet from the centrifugation (which contains the insoluble proteins), 150 µl 100 mM Tris pH 7.4, 150 mM NaCl, 8M urea was added and incubated at 4°C for 20 min. The urea buffer was prepared fresh and used within a week. Both the fractions were bound to Ni-NTA beads and purified as described for anti-FLAG M2 binding.

### Mammalian cell experiments

HEK293T cells were cultured at 37°C in DMEM supplemented with sodium pyruvate. Upon reaching 70% confluency in a 6-well plate, cells were transfected with individual plasmids by lipofection (TransIT®-293, Mirus Bio). 48 hr after transfection, the media was removed and the cells were washed with PBS, harvested by trypsinization, pooled into one 15 ml Falcon tube and harvested by centrifugation. Cells were washed with 2 ml PBS and once with 4 ml PBS to remove any untransfected plasmids and flash frozen with liquid nitrogen until use.

450 µl Xtractor buffer was added to the frozen cell pellet with 0.1 mM Pefabloc, 3 µl murine RNase inhibitor and 40 µl 100 mM stem-loop RNA. Cells were incubated for 20 min and centrifuged at 21,000 g for 30 min. The clarified lysate was bound to magnetic anti-FLAG M2 beads as described for the anti-FLAG M2 binding assay.

### Data analysis

#### PacBio long read data

Association of variants to unique barcodes (PacBio data) was performed using a custom script (see Supplementary data). Only reads with 10 or more counts were used for further analysis. Filtering was used to ensure only non-duplicated barcodes were used for further analysis.

#### Illumina short read data

Reads were converted from BCL to FASTQ using bcl2fastq. Paried reads were joined using PANDAseq. Reads for each barcode were counted in both input and output (cDNA) sample. Only barcodes that were sequenced at least 5 times in each replicate and were present in all input and output (cDNA) samples were used for further analysis. To calculate enrichment scores, counts for each barcode were normalized by the total reads in their respective sample. We then employed two steps to generate a final enrichment score. First, we calculated a log_2_ ratio of output reads by input reads. Second, for each variant we subtracted the WT log_2_ score. In the case of the protease substrate specificity assay, the enrichment score was inverted by multiplying the dataset by –1. For protein solubility analysis, a log_2_ ratio of soluble to pellet (insoluble) fraction was used to generate solubility scores.

### Statistical tests

Proportion tests were performed using the prop.test function from the statsmodels package in Python 3.0, executed within a Google Colab environment. Specifically, the Z-test for proportions was used to assess statistical significance.

### Accession numbers

Barcode sequencing reads were deposited in the National Center for Biotechnology Information (NCBI) Sequence Read Archive under the BioProject accession PRJNA1329341.

## Supporting information

Supplementary results

## Contributions

Y.A and S.F conceived the project. Y.A designed and performed experiments and analysis. S.F supervised the research. K.L.B, S.G and C.O contributed codes for analysis. C.Q. and J.C provided advice on general strategy and data analysis. Y.A and S.F wrote the paper with input from all authors. Y.A and S.F are in the process of filing a patent application.

## Acknowledgements

We thank Miklos Sahin-toh, Karl Barber and Stephen Elledge for sharing data, members of the Fields and Queitsch-Cuperus labs for advice, and Anna Jansen, Elliott Burke and Ruth Groza for their work in the early stages of the project. We thank Tobias Jores, Douglas Fowler and Eric Phizicky for comments on the manuscript. This work was supported by RM1HG010461 and R21HG013504 from the National Human Genome Research Institute of the NIH.

