## Supplementary results for "Dosa: A method to covalently barcode proteins for high throughput biochemistry"

**for high throughput biochemistry**

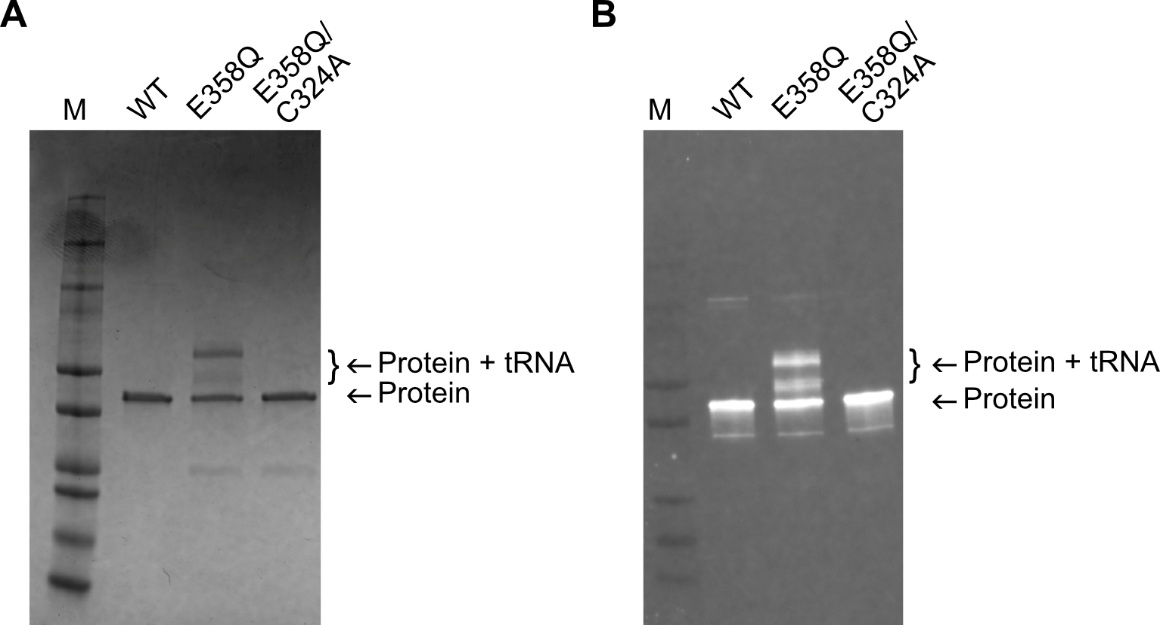

**Fig. S1. *In vivo* covalent linking of TrmA (E358Q) mutant to tRNA.**  **A.** SDS-PAGE stained with Coomassie Blue showing purified TrmA wild type (WT) and two TrmA mutant proteins. The TrmA (E358Q) mutant exhibits an additional band corresponding to a linked tRNA species, which is absent in the WT and the catalytically dead double mutant (E358Q, C324A). **B.** Western blot analysis of the same samples shown in **A**, probed with an anti-6X His antibody, confirming the presence of the TrmA-tRNA covalently linked species of the TrmA (E358Q) mutant. The two bands of “Protein + tRNA” in the E358Q mutant are likely due to tRNA heterogeneity.

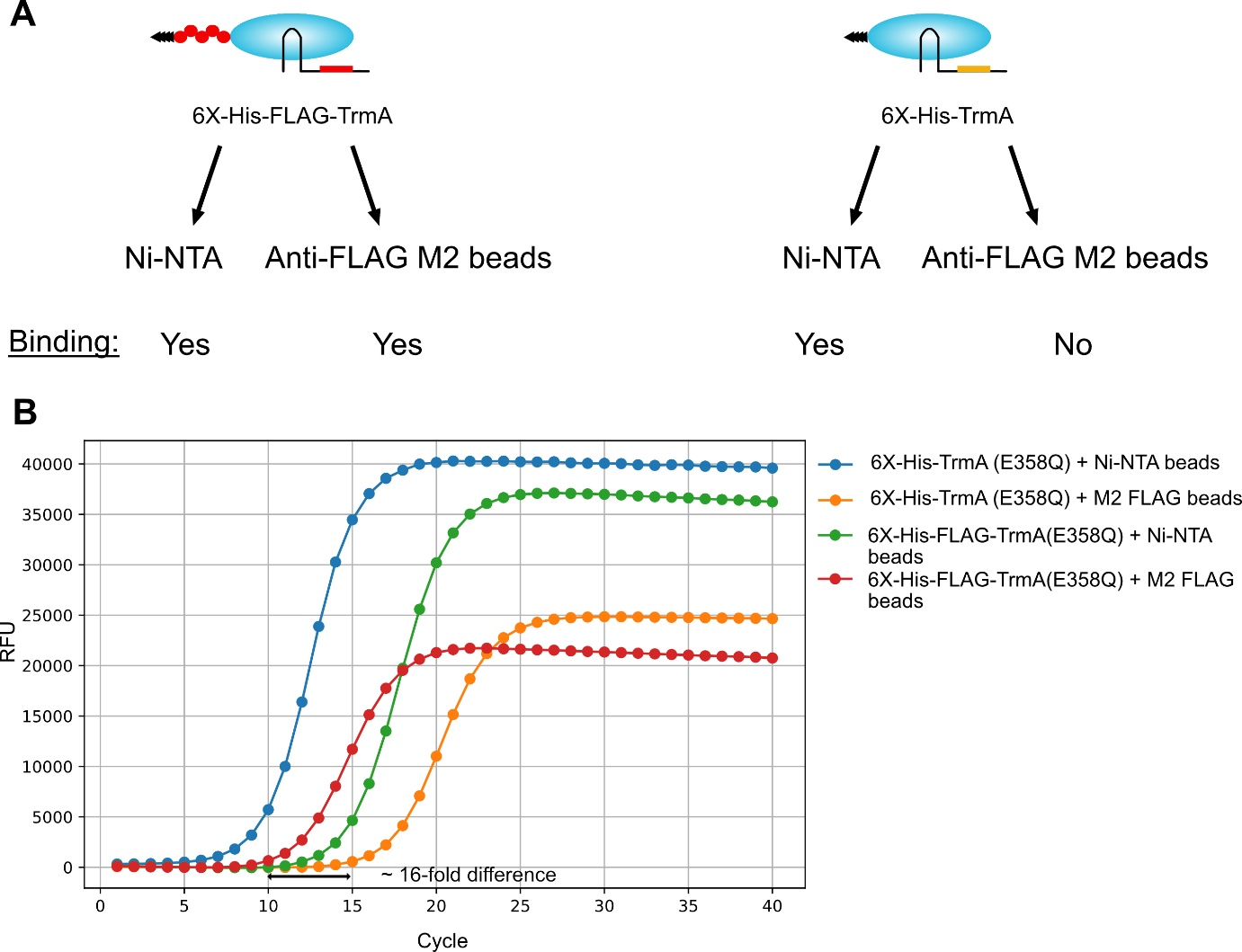

**Fig. S2.** **Assay of FLAG epitope binding to M2 anti-FLAG antibody**. Lysates from cells expressing the indicated constructs were bound to beads containing either Ni-NTA or M2 anti-FLAG antibody, and the bound barcoded proteins were reverse transcribed and amplified by qPCR. Both 6X-His-TrmA (E358Q) and 6X-His-FLAG-TrmA (E358Q) barcodes could be amplified using Ni-NTA beads. The 6X-His-FLAG-TrmA (E358Q) RNA barcode was selectively enriched by ~16-fold on anti-FLAG M2 beads. The 16-fold difference was calculated from quantification threshold (Cq).

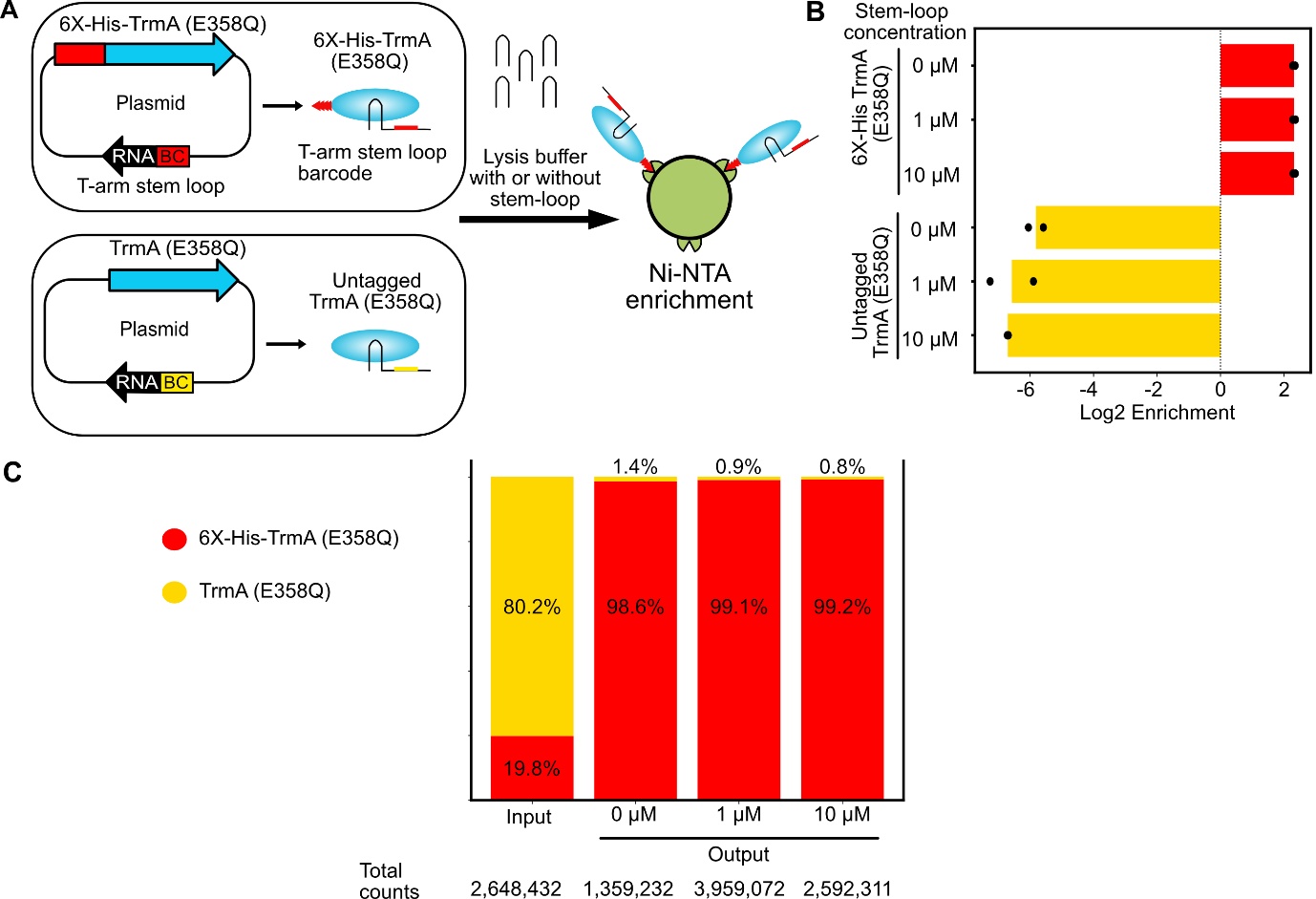

**Fig.S3. Preventing barcode mixing after cell lysis by blocking TrmA (E358Q) with the tRNA stem-loop that contains no barcode. A.** Two populations of *E. coli,* one encoding 6X-His-TrmA (E358Q) and another encoding untagged TrmA (E358Q) were mixed at equal density and induced for production of the barcoded protein. Cells were lysed in the presence of either 0, 1 or 10 µM of the 19-base WT stem-loop RNA and the lysate was enriched on Ni-NTA beads, followed by reverse transcription, PCR and sequencing. Plasmids isolated from an identical induced culture was used as input for sequencing. **B.** Enrichment plots from the experiment in **A** shows enrichment of the barcode specific for 6X-His-TrmA (E358Q) and depletion of the barcode corresponding to untagged TrmA (E358Q). The specific enrichment (red bars) does not significantly increase upon increasing stem-loop concentration (prop test: p-value <10⁻²⁰), but the proportion of the non-specific barcode (yellow bars) reduces with increasing stem loop concentration (n=2). **C.** Stacked bar plots showing proportions of barcodes in the input and output selections at varying stem-loop concentrations.

**Supplementary discussion for Fig. S3.**

We sought to investigate whether non-cognate barcodes, liberated from neighboring cells during cell lysis, could link to non-orthogonal TrmA (E358Q) fusion proteins that had not yet formed a link. To address this possibility, we pooled cells containing two populations of plasmids: 6X-His-TrmA (E358Q) with one barcode, and untagged TrmA (E358Q) with a different barcode. After cell lysis, upon enrichment of the 6X-His-TrmA (E358Q) on Ni-NTA beads, only the barcode corresponding to the tagged protein should be enriched. However, non-orthogonal linkage of barcodes during lysis would lead the barcode associated with the untagged TrmA (E358Q) to also be enriched. Plasmid DNA isolated from the pooled culture was used as the input, and the barcodes enriched on Ni-NTA beads were used to calculate output proportions. The average ratio of barcodes corresponding to 6X-His-TrmA (E358Q) in the input pool was 0.24 and after selection on Ni-NTA beads increased to 70 in output pool (prop test: p-value < 10⁻²⁰), corresponding to 293-fold enrichment (ratio of output [6X-His-TrmA (E358Q)/TrmA (E358Q)] / input [6X-His-TrmA (E358Q)/TrmA (E358Q)]) when no stem loop was added. During cell lysis, we added a 19-base stem-loop at two concentrations (1 and 10 µM) to quench any unreacted TrmA (E358Q) binding sites. Our hypothesis was that addition of a stem-loop sequence would reduce any barcode swapping if it occurred. Since the 19-base stem loop does not contain a barcode sequence, it would be silent in the sequencing analysis. The results indicate that the addition of the stem-loop did not result in any increase in specific barcode [6X His-TrmA (E358Q)] enrichment but it led to a depletion of ~1.9-fold of the non-orthogonal barcode [untagged TrmA (E358Q)] at the higher stem-loop concentration (Fig. S3B,C). These results indicate the stem-loop may prevent non-specific binding of the RNA barcode to proteins.

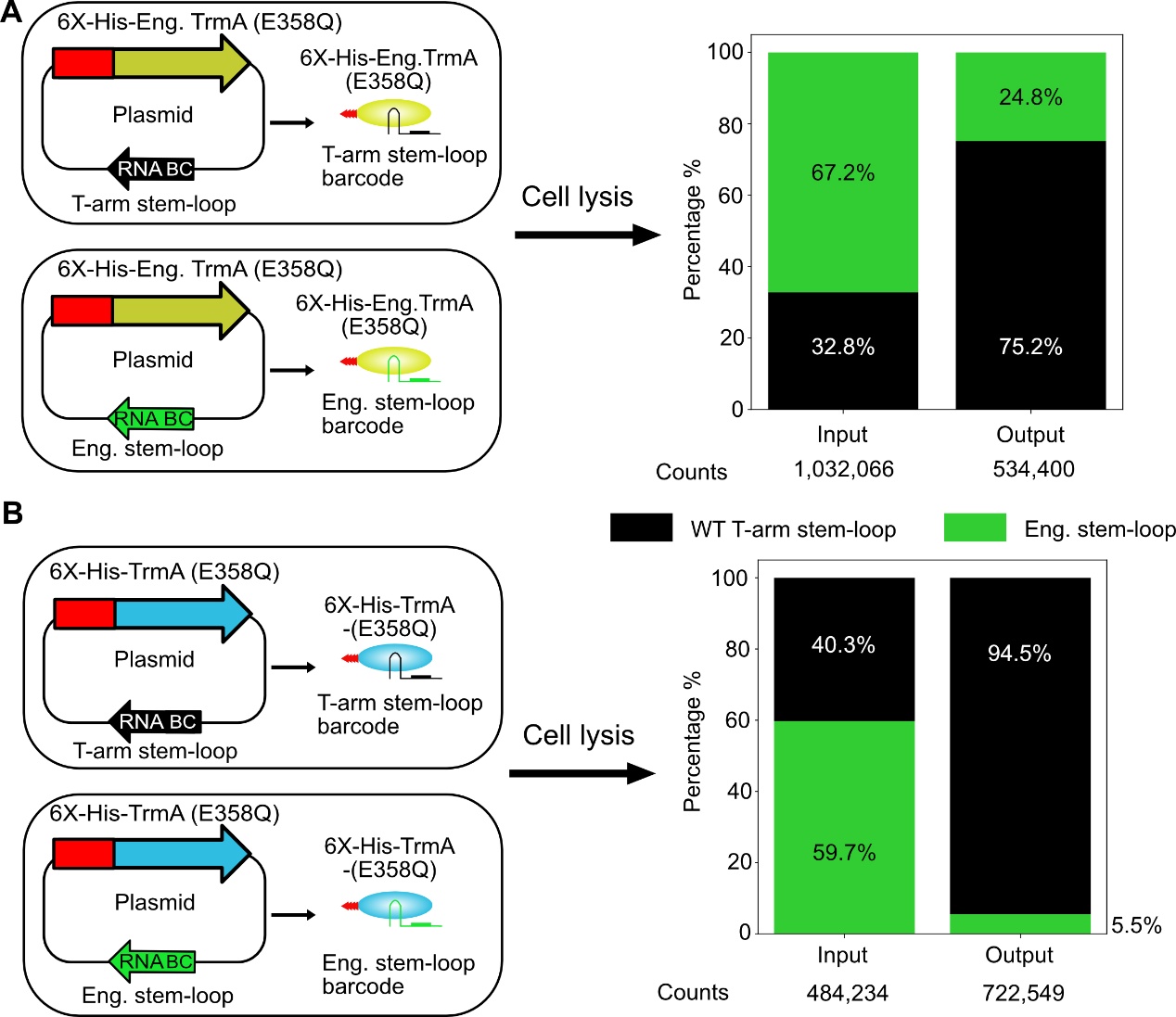

**Fig. S4. Testing the orthogonality of an engineered TrmA mutant with an engineered stem-loop mutant**. **A.** Two populations of *E. coli* containing a plasmid encoding 6X-His-engineered TrmA (E49R, R51E, E358Q) and either the WT *E. coli* T-arm stem-loop (5’GCTGTGTTCGATCCACAGC 3’) or an engineered stem-loop (5’GCTGTGTT*A*G*GT*TCACAGC 3’, mutations are underlined). Each population contained a unique barcode. After cell lysis, the 6X-His-TrmA (E358Q) was bound to Ni-NTA beads for enrichment. The bound barcodes were used as output and the plasmid samples isolated from induced cultures was used as input. **B.** Two populations of *E. coli* containing a plasmid encoding 6X-His-TrmA (E358Q) and either the WT *E. coli* T-arm stem-loop (5’ GCTGTGTTCGATCCACAGC 3’) or an engineered stem-loop (5’ GCTGTGTT*A*G*G*T*T*CACAGC 3’, mutations underlined). Each population contained a unique barcode. Lysates were treated as in **A**.

**Supplementary discussion for Fig. S4**

Full-length tRNA exhibits approximately ten times greater affinity for TrmA than the T-arm stem-loop^1^. Given that cellular concentrations of tRNA are much higher than those of the recombinant stem-loop barcode sequence and the efficient linking between tRNA and TrmA (E358Q) (Fig. S1), another group has engineered the TrmA protein to link to an orthogonal T-arm^1^. We refer to these components as engineered TrmA (E49R, R51E, E358Q) and engineered stem-loop, which differs from the WT stem loop by 3 bases (Fig. S4). We tested whether 6X-His-engineered TrmA preferentially links to the WT stem-loop or the engineered stem-loop (Fig. S4A) in an *in vivo* context using a pool of *E. coli* culture that contains one of the stem-loop sequences with its unique barcode. As the stem-loop RNA barcode sequence is expressed under the same promoter and terminator, we reasoned that the RNA barcode expression level would reflect the plasmid proportions. Therefore, we used the plasmids from the pooled culture as an input for sequencing. Cell lysates were bound to Ni-NTA beads to enrich barcoded proteins as the output for sequencing. The ratio of the WT stem-loop barcode to the engineered stem-loop barcode in the input was 0.5 and increased to 3 in the output after selection, corresponding to a ~6-fold enrichment of WT stem-loop barcode (ratio of output [WT stem-loop barcode/engineered stem-loop barcode] /  input [WT stem-loop barcode/engineered stem-loop barcode]). A similar analysis with TrmA (E358Q) and the WT stem-loop vs. engineered stem-loop showed that TrmA (E358Q) robustly linked to the WT stem-loop (Fig. S4B). The ratio of the WT stem-loop barcode to the engineered stem-loop in the input was 0.6 and increased to 17 in the output pool after selection, corresponding to a ~29-fold enrichment of the WT stem-loop barcode (ratio of output [WT stem-loop barcode/engineered stem-loop barcode] /  input [WT stem-loop barcode/engineered stem-loop barcode]).These findings prompted us to use TrmA (E358Q) rather than the engineered version for further studies. Although our results do not correspond with the previous study^1^, our experiments carried out the RNA linking *in vivo*, while the original study used *in vitro* experiments.

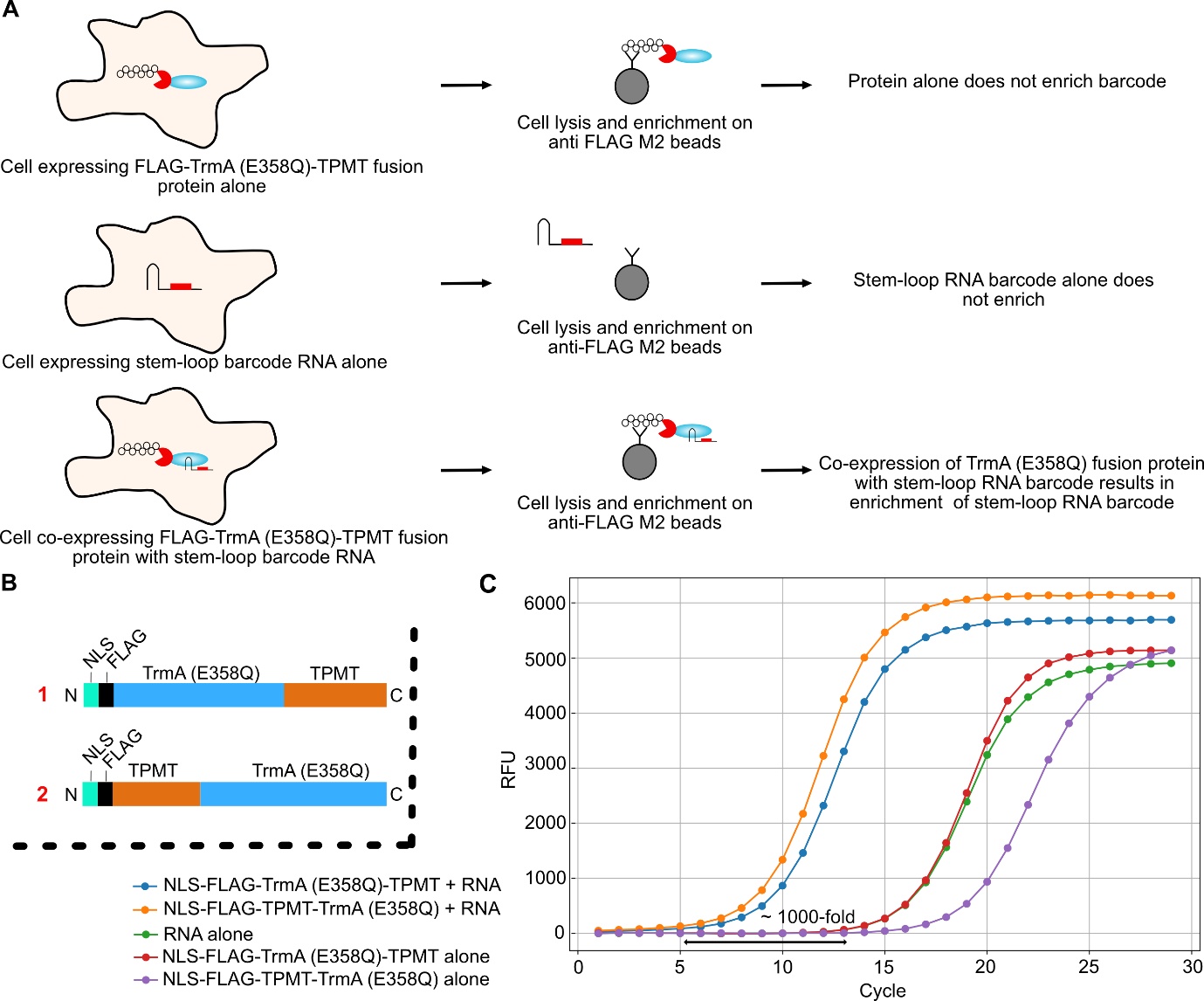

**Fig. S5. Optimization of the Dosa method in mammalian cells.** **A.** Scheme for the experiment. Cells were transfected with either FLAG-TrmA (E358Q)-TPMT fusions alone or RNA barcode alone or co-transfected with both FLAG-TrmA (E358Q)-TPMT fusion and the RNA barcode. Cells were lysed and processed independently for anti-FLAG M2 binding and enrichment was monitored by qPCR. **B.** Constructs used in the experiment are shown. NLS, Nuclear localization sequence. **C.** Representative qPCR amplification curves of the indicated constructs after their enrichment on anti-FLAG M2 beads (n=2). Only constructs expressing both the TrmA (E358Q) fusion protein and the RNA barcode are enriched in the qPCR. The difference shown was calculated from quantification threshold (Cq).

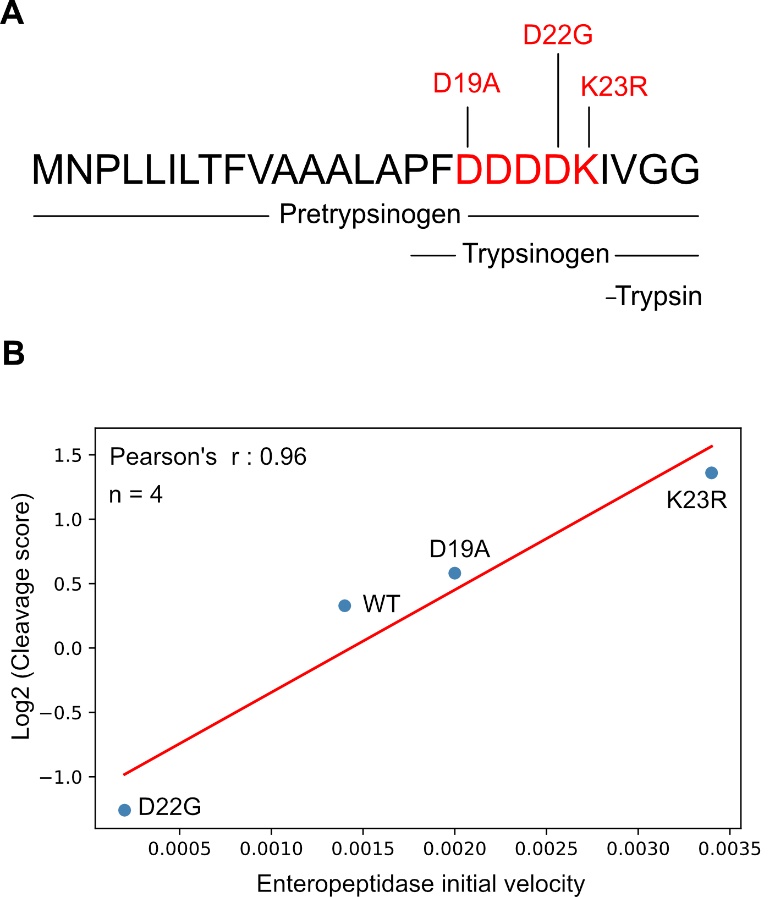

**Fig. S6. Enteropeptidase cleavage and correlation. A.** Sequence of human cationic trypsinogen (Uniprot: P07477). Pretrypsinogen is cleaved co-translationally into trypsinogen by endoplasmic reticulum proteases. Enteropeptidase acts on the trypsinogen cleavage peptide (DDDDK, highlighted in red) to produce active trypsin. The C-terminus of the FLAG epitope is identical to the trypsinogen cleavage peptide (FLAG epitope: DYKDDDDK). Clinical mutations in the trypsinogen gene are highlighted above the sequence. **B.** Pearson correlation between log₂-transformed cleavage scores from Dosa and initial velocity measurements of enteropeptidase for the indicated variants^2,3^. Red line indicates linear regression fit.

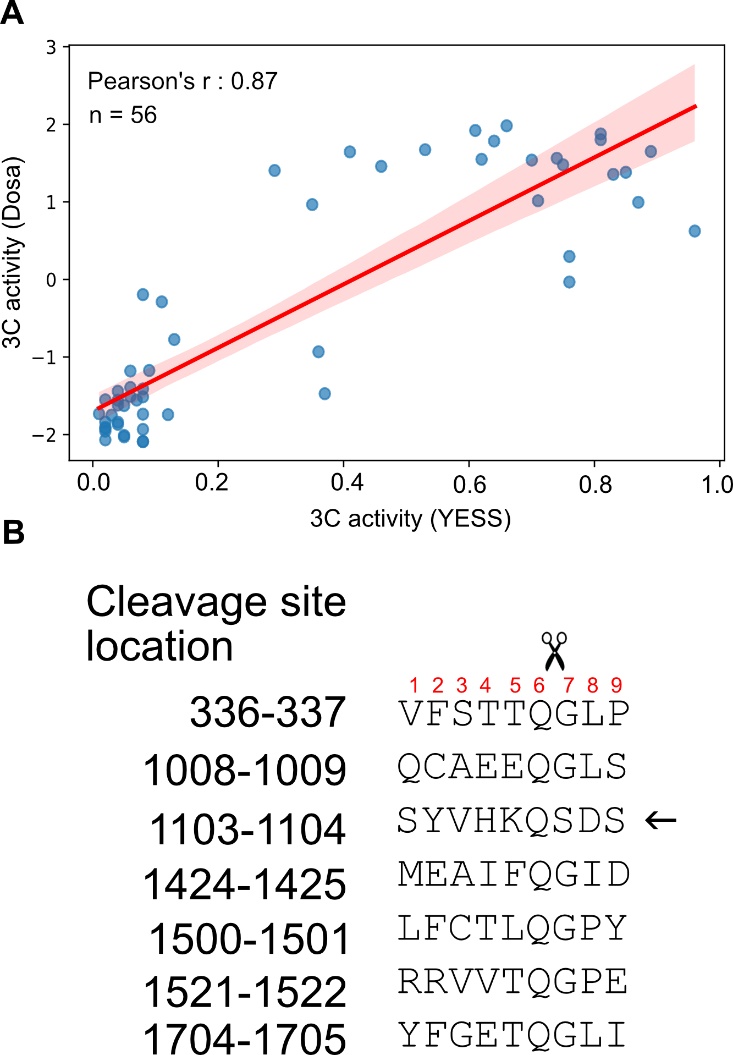

**Fig. S7. HRV 3C cleavage and correlation. A.** Pearson’s correlation between Yeast Endoplasmic Reticulum Sequestration Screening (YESS) of 3C protease on substrate variants and Dosa-based 3C protease cleavage assay. Only data for positions 6, 7 and 8 are available^4^. Red line indicates linear regression fit. **B.** Alignment of HRV 14 polyprotein, highlighting cleavage positions. Scissors indicate the position of the bond that is cleaved, with the serine in position 7 indicated by an arrow. (Sequence from Uniprot: P07210).

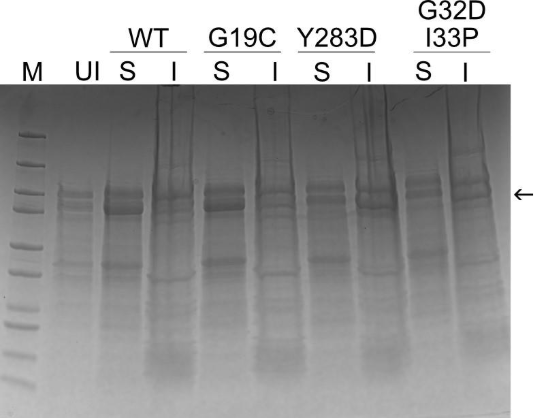

**Fig. S8.** **Solubility analysis of Maltose Binding Protein (MBP)-TrmA (E358Q) fusion mutants using SDS-PAGE.** Arrow indicates the MBP-TrmA (E358Q) band. The G19C mutant shows comparable solubility to the WT, whereas the Y293D mutant and G32D, I33P double mutant are insoluble. Cell lysates were centrifuged to separate soluble and insoluble fractions. The pellet fraction contains insoluble proteins. M, Protein marker; UI, Uninduced; S, Supernatant; I, Insoluble fraction.

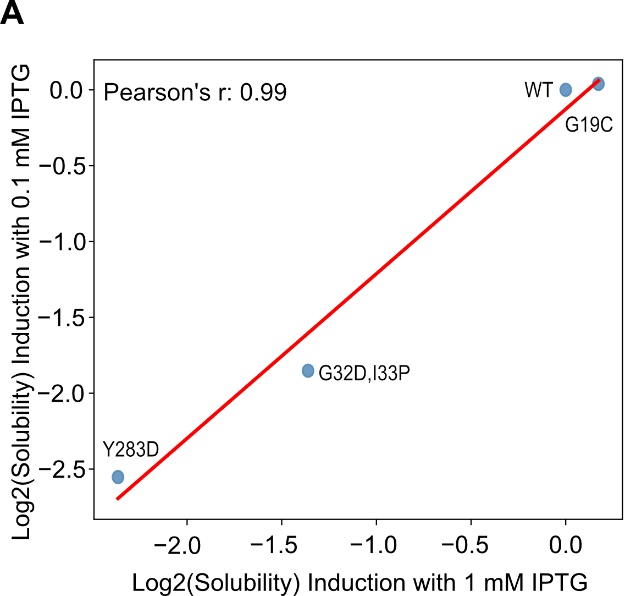

**Fig. S9.**  **Correlation plots for solubility analysis of MBP variants. A.** Pearson’s correlation plot of Log_2_ Solubility scores for MBP variants expressed with 0.1 mM IPTG and 1 mM IPTG using the Dosa assay (n=2). Red line indicates linear regression fit. In *E. coli*, inducer concentration and temperature determine the outcome of the recombinant protein solubility. Overexpression of proteins saturates the capacity of cellular protein folding machinery leading to production of insoluble proteins. To ensure that Y283D and G32D/I33P variant are bonafide insoluble variants, we tested a lower induction concentration (0.1 mM IPTG). These results show that induction at 1 and 0.1 mM IPTG give the same results.

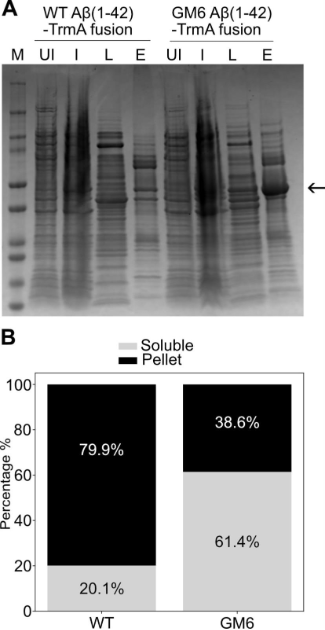

**Fig. S10. Amyloid Aβ(1-42)-TrmA (E358Q) model selection and solubility.** **A.** Small-scale purification of Aβ(1-42)-TrmA (E358Q) fusion protein variants. M, Protein marker; UI, Uninduced sample; I, Induced sample; L, Crude lysate; E, elution from Ni-NTA beads (indicating soluble fraction). The arrow indicates the position of the expected band. The GM6 Aβ(1-42)-TrmA (E358Q) fusion is more soluble than the WT Aβ(1-42)-TrmA (E358Q) fusion as indicated by more protein in the elution than WT. **B.** The solubility of WT and GM6 Aβ(1-42)-TrmA (E358Q) fusions determined by Dosa (stacked bar plots show mean of n=2).

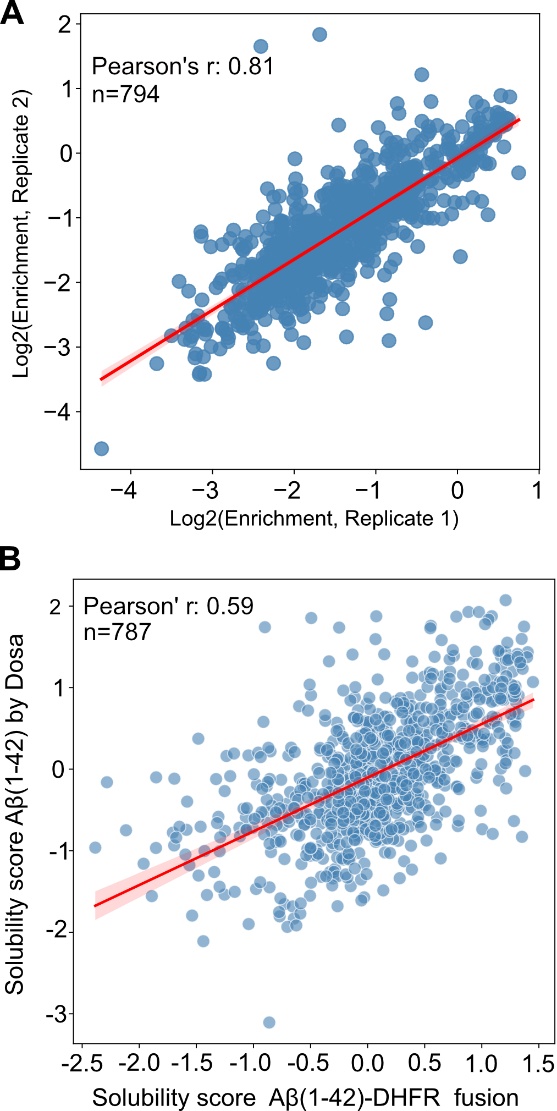

**Fig. S11. Correlation plots for Aβ(1-42) datasets.  A.** Pearson’s correlation between two replicates from this study. **B.** Pearson’s correlation between solubility scores from Dosa and from experiments using an Aβ(1-42)-DHFR fusion^5^. Red line indicates linear regression fit in both plots.

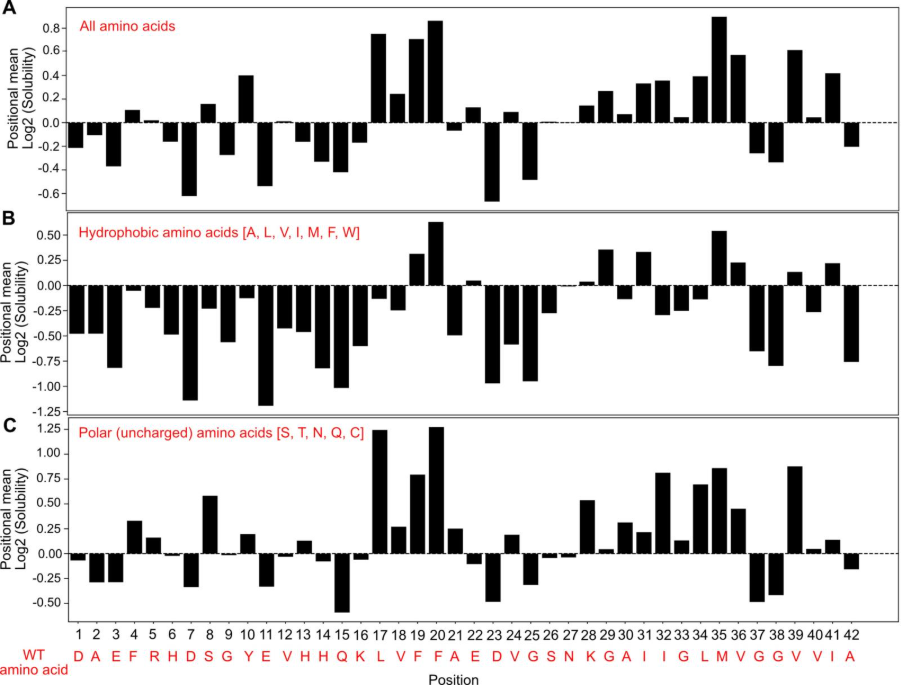

**Fig. S12. Positional means of amino acid substitutions in Aβ(1-42) solubility assay. A.** Positional mean at each position of all amino acid substitutions. **B.** Positional means of only hydrophobic amino acid substitutions (amino acids A, L, V, I, M, F, W). **C.** Positional means of only polar amino acid substitutions (amino acids S, T, N, Q, C).

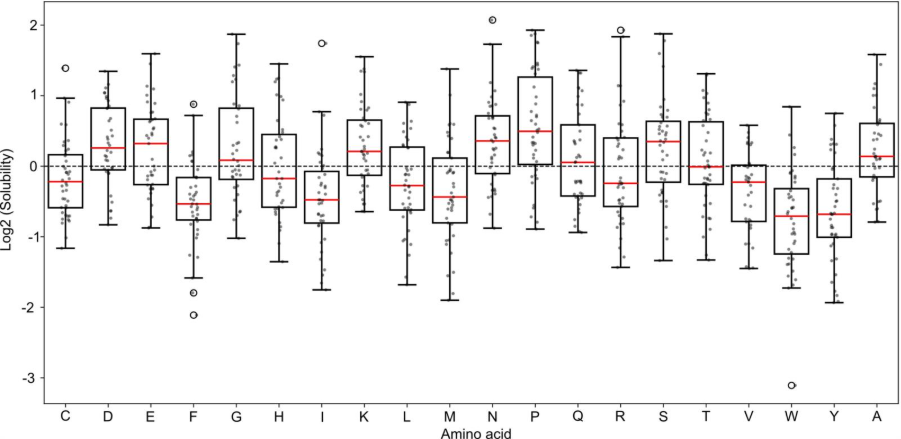

**Fig. S13. Effect of each amino acid substitution on Aβ(1–42) aggregation.** 
Variants are grouped by the identity of the substituted amino acid. Red line indicates the median of all datapoints for each amino acid. Each point represents a single-substitution variant.

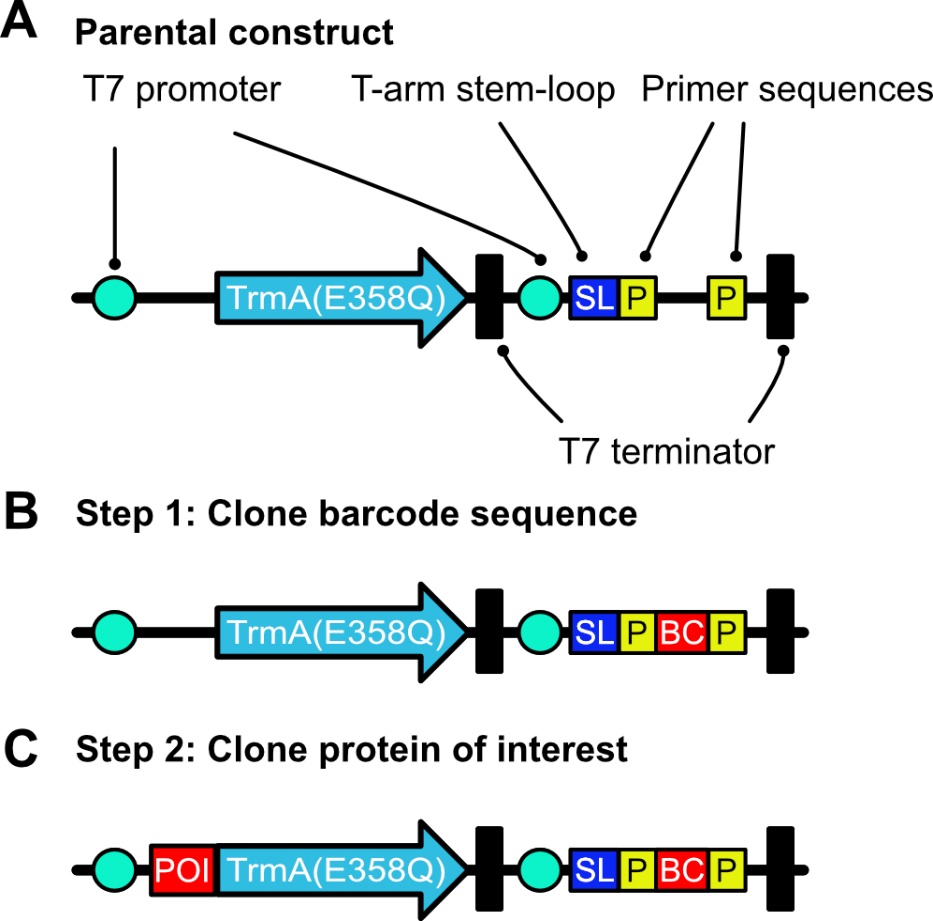

**Fig. S14. Construction of plasmid libraries.** **A.** Parental construct devoid of Golden Gate restriction enzyme sites Esp3I and BsaI. TrmA (E358Q) is expressed from a T7 promoter and a T7 terminator. The RNA for barcoding is expressed as a separate transcription unit. The RNA has the T-arm stem-loop and primer sequences that allow the RNA barcode to be reverse transcribed and amplified for sequencing. **B.** Random barcode (BC) sequences are inserted into the plasmid by digesting and ligating the plasmids with Esp3I. **C.** Pre-barcoded plasmid pool obtained from step 1 is used for insertion of the protein of interest (POI) using BsaI digestion and ligation.
